# Design of an intracellular aptamer-based fluorescent biosensor to track burden in E. coli

**DOI:** 10.1101/2024.09.23.614511

**Authors:** Alice Grob, Tom Copeman, Sifeng Chen, Yuval Elani, Elisa Franco, Francesca Ceroni

## Abstract

Cell burden impacts the performance of engineered synthetic systems. For this reason, there is great interest toward the development of tools to track burden and improve biotechnology applications. Fluorogenic RNA aptamers are excellent candidates for live monitoring of burden because their production is expected to impose a negligible load on transcription resources. Here we characterise the performance of a library of aptamers when expressed from different promoters in *E. coli*. We find that aptamer relative performance is dependent on the promoter and the strain, and that, contrary to expectation, aptamer expression impacts host fitness. By selecting two of the aptamers with brighter output and lower impact, we then design an intracellular biosensor able to report on the activation of the burden response in engineered cells. The sensor developed here adds to the collection of tools available for burden mitigation and may support bioprocessing applications where improved host performance is sought.

## Introduction

In engineered cellular hosts, synthetic and native gene expression processes compete for the cellular resources available within the cell, resulting in cellular burden^1^. Burden is a response to cellular overload, known to affect both cell growth and the functionality of heterologous constructs. Extensive work has been carried out to characterise and mitigate burden in bacteria, to identify strategies and approaches that improve the performance of synthetic genetic constructs and their downstream applications in therapy and biomanufacturing^2^. We previously investigated global transcriptional changes occurring in *E. coli* when burden is triggered by a library of genetic constructs^3^. By RNAseq experiments, we found that activation of the native htpG1 promoter, measured as abundance of resulting mRNA present within the cells overtime, occurs already at 15 minutes after induction of heterologous protein expression. We used this promoter to design a biomolecular feedback controller capable of mitigating burden and improving expression yields over long-term cultivations. While the feedback could sense and respond to burden, it did not provide live visualisation of the burden imposed by a selected genetic construct. This feature would be of significant help for researchers seeking to quantify and track burden as a function of time in living cells, providing the ability to select for low footprint variants which are better performing in each context.

Here, we use htpG1 to develop RNA-based biosensors for the live dynamic tracking of the burden response in growing engineered *E. coli* populations. As reporters, our biosensors use fluorogenic RNA aptamers, short hairpin forming RNAs that bind to small fluorophore molecules (dye) resulting in fluorescence^4-6^. Many fluorogenic aptamers have been developed, to mitigate issues related to dim signal intensity and instability. Among these, variants of Spinach^7-9^, Corn^10^, Broccoli^11,12^, Beetroot^13^, Mango^14^, Malachite Green^15^, and Pepper^16,17^ were developed widening the applicability of aptamers as synthetic biology tools^118-21^. Fluorogenic RNA aptamers represent a step forward when compared to other RNA detection techniques, as they allow tracking and direct quantification of RNA expression in living cells^22^. Another advantage is the easy cloning required for aptamer insertion in any synthetic RNA of interest^14^ which makes them suitable for high throughput and dynamic screening of RNA expression and biosensing in a wide range of biotechnology applications, including the design of complex synthetic biology circuitry like RNA-based toggle switches^24^ and biosensors^25-27^, mRNA^23^ or ribosome tagging^28^ and mRNA tracking^29, 30^ in a variety of hosts^10, 16, 17, 22, 31^. For some of the aptamers, like Pepper, adapting the small HBC fluorophore molecule used in combination with the aptamer, also generates a rainbow of spectral fluorescence from cyan to red, enabling selection of the spectrum of choice and reducing the need for further cloning when a non-compatible fluorescent reporter is expressed. Finally, RNA aptamers allow for quicker expression detection compared to protein reporters which possess longer folding and maturation times^16^.

While fluorogenic aptamers offer many advantages as the readout of our biosensors, and a multitude of aptamer variants is available for RNA-based cell engineering and studies, these variants have been characterised in different hosts, making it difficult to compare their performance and to select the best aptamer for a particular application. Aptamers are also most often expressed from strong viral promoters and wider characterisation of their behaviour is needed to identify suitable ones for tracking live expression from weaker promoters, like the native htpG1.

To address these challenges, here we characterise a library of seven fluorogenic RNA aptamers in two *E. coli* strains, investigating their relative performance when expressed by different synthetic and native promoters. We observed how the aptamer relative performance is dependent on the promoter and the cellular host of choice and compared this with *in vitro* performance in commercially available cell free protein expression systems made of both purified transcription/translation components and cell lysate buffers. We also find that certain fluorogenic aptamers can impact host physiology and lead to growth defects, and that their impact appears to be correlated to the complexity of their scaffold. This surprising result is a call for caution when selecting biosensors for a given application, especially when cell-based work is envisioned. Finally, by adopting the aptamer displaying the brightest signal across contexts together with the lowest impact on the host, we designed an intracellular burden biosensor able to report on the activation of *E. coli* burden response, thus expanding the applications of fluorogenic aptamers towards biosensing of native physiological processes and downstream biotechnology applications.

## Results

To select a suitable readout for the design and assembly of our burden biosensor, we started by characterising the performance of a library of fluorogenic RNA aptamers when these are expressed from diverse types of promoters. We selected a library (Figure 1a) of native and synthetic bacterial promoters of different strength (a weaker and stronger variant of pBAD^32^, pLux and J23118 from the Anderson promoter library) comparing aptamer expression profiles to expression when the T7 viral promoter is adopted. Among the available aptamers, we selected seven aptamers of recent development; five representing variations of the Pepper aptamer (Pep) (F30-1xPepper, F30-4xPepper, F30-8xPepper^16^, tRNA-1xPepper^16^, tRNA-DF30-(2x)Pepper^17^) and two variations of the Broccoli aptamer (Broc) (F30-4xBroccoli^11^ and tRNA-enhanced-Broccoli^33^), for which characterisation is currently available mainly in mammalian cells or in *in vitro* studies. To allow comparison of aptamer performance across conditions, we benchmarked their fluorescence and dynamics against the standard tRNA-Broccoli aptamer.

**Figure 1.**
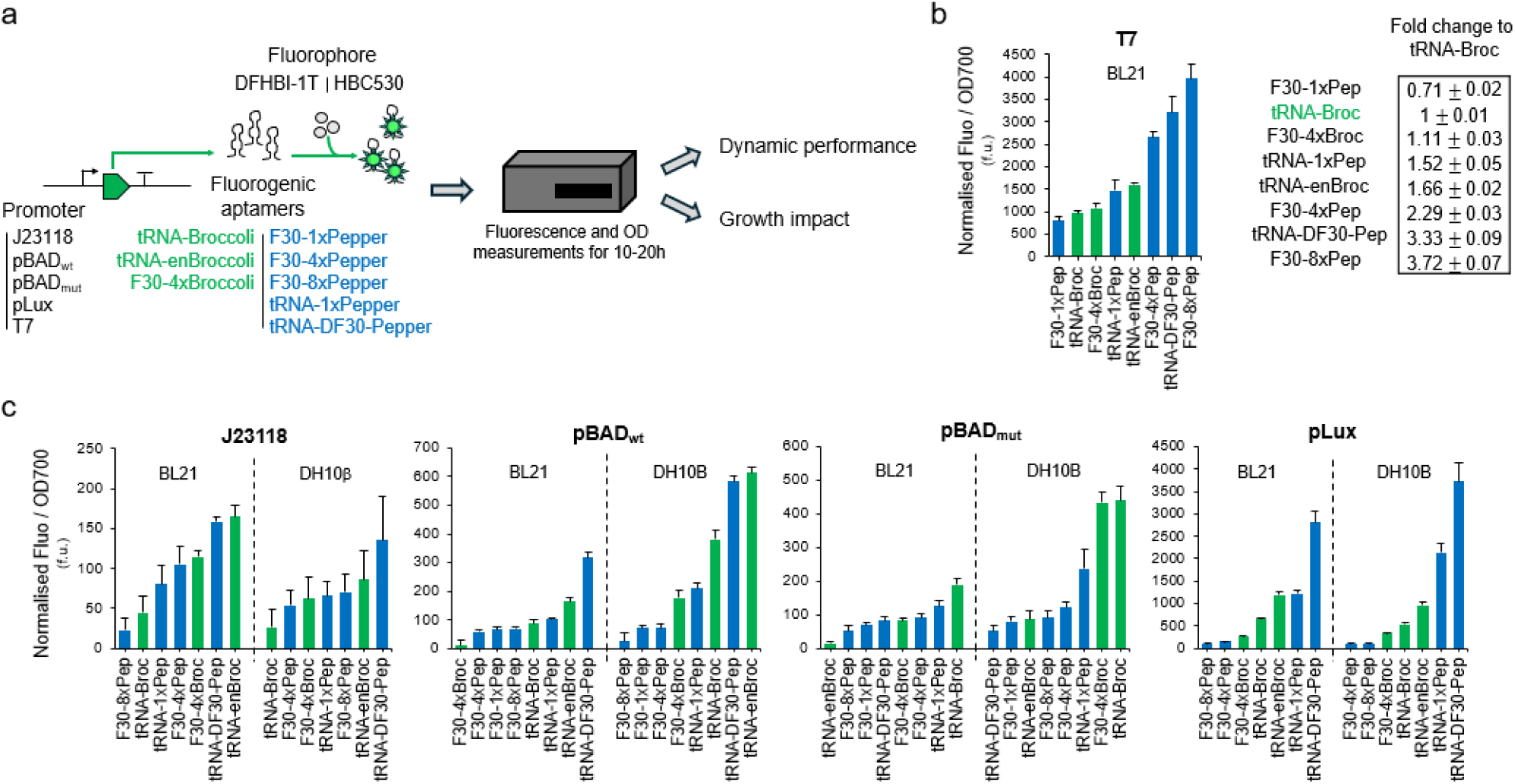
Context dependency performance of Pepper and Broccoli aptamers when expressed in *E. coli*. **a)** Schematic of plate reader-based workflow used to characterise Pep and Broc expression in *E. coli*. A small library was generated combining five promoters (J23118, pBAD_wt_, pBAD_mut_, pLux and T7) with three Broc aptamers (tRNA-Broc, tRNA-enBroc and F30-4xBroc; green) and five Pep aptamers (F30-1xPep, F30-4xPep, F30-8xPep, tRNA-1xPep and tRNA-DF30-Pep; blue). This library was transformed into *E. coli* BL21 and DH10B when appropriate. Transformed cells were cultured overnight in M9 medium and diluted the following morning to OD700∼0.05. Following induction, samples were loaded into a 96-well plate and fluorophore molecules, 160uM DFHBI-1T or 2uM HBC530, were added to samples containing respectively Broc or Pep aptamers. Subsequently, fluorescence levels and OD700 were measured to assess both dynamic performance and growth. **b)** Expression levels of T7 constructs in BL21 following induction with 1mM IPTG. Bar chart represents fluorescence level per OD700 of Broc aptamers (green) and Pep aptamers (blue) at maximum growth rate, normalised by subtracting that of induced samples containing no fluorophore. Data are presented as mean values + SEM. Table indicates the resulting fold change +/- SEM of fluorescence in comparison to tRNA-Broc highlighted in green. (n=4) **c)** Expression levels of J23118, pBAD_wt_, pBAD_mut_ and pLux constructs in *E. coli* BL21 and DH10B with bar charts of fluorescence level per OD700 of Broc aptamers (green) and Pep aptamers (blue) at maximum growth rate, normalised by subtracting that of induced samples containing no fluorophore. Expression of pBAD_wt_, pBAD_mut_ constructs is induced with 0.2% arabinose, while expression of pLux constructs is induced by 10uM HSL. Data are presented as mean values + SEM. (n=4)

To enable high throughput characterisation of steady state and dynamic aptamer expression levels, we set up a plate reader-based workflow (Figure 1a). Cells were cultured overnight in M9 medium and diluted back to the beginning of exponential phase (optical density, OD700 ∼ 0.05) the following morning. Samples were then induced, and fluorescence assessed over a 10 hour-window. Fluorescence per OD700 was normalised to that of induced cells not supplied with fluorophore molecule. Plate reader measurements allowed us to also gain information on the impact of these aptamers on the host, quantified as changes in OD700, something that, to the best of our knowledge, has not been previously reported.

We first considered intensity levels of the aptamers when they are expressed from the T7 promoter in BL21 bacterial cells. At maximum growth rate post induction, F30-8xPep, tRNA-DF30-Pep and F30-4xPep displayed the highest normalised fluorescence per OD, all of them being more fluorescent than the Broccoli variants and resulting, respectively, in 3.72-, 3.33- and 2.29-fold increase in fluorescence when compared to tRNA-Broc chosen as the reference for relative aptamer strength. Remaining tRNA-enBroc, tRNA-1xPep, F30-4xBroc and F30-1xPep aptamers yielded 1.66, 1.52, 1.11 and 0.71 fluorescence fold change, respectively (Figure 1b).

We then studied the influence of promoter strength on aptamer fluorescence. We expressed aptamers in BL21 and DH10B from the weaker and stronger variants of the pBAD promoter, pLux and J23118, expecting that the relative ranking of aptamer strength would be conserved across promoters (Figure 1c, Figure S1). However, we noticed that aptamers strength is affected by the context of expression, i.e. the promoter and the bacterial strain. F30-8xPep did not appear to be the brightest aptamer when expressed from these bacterial promoters (Figure 1c and Figure S1). Also, while in both BL21 and DH10B, tRNA-Broc displayed the lower fluorescence (lower in DH10B and second to lower in BL21 after F30-8xPep) when expressed under J23118, for the other promoters in the same strains tRNA-Broc resulted in medium to high fluorescence compared to the other aptamers, highlighting the impact of the genetic context and expression host on aptamer performance (Figure 1c). In BL21, tRNA-DF30-Pep and tRNA-enBroc resulted in higher expression (3.54- and 3.68-fold change, respectively) compared to tRNA-Broc when J23118 was used (Figure S1a). In DH10B, the same aptamers yielded 3.22- and 5-fold change, respectively. When pBAD_wt_ was adopted, tRNA-DF30-Pep and tRNA-enBroc were also the brightest aptamers, resulting in 4.27- and 1.87-fold change in BL21 cells, and 1.57- and 1.61-fold change in DH10B cells, respectively (Figure S1b). Finally, with pLux, the highest fluorescence performance was achieved with all aptamers compared to other promoters, with tRNA-DF30-Pep, tRNA-1xPep and tRNA-enBroc displaying the highest output of respectively 4.73-, 1.87- and 1.81-fold change in BL21 cells, and 7.03-, 4.03- and 1.87-fold change in DH10B cells (Figure S1c). An exception was observed for pBAD_mut_, for which tRNA-Broc resulted in the highest output in both strains when compared to the other aptamers (Figure S1d).

Looking then at the dynamic response of the aptamers expressed from T7 over 10 hours, we noticed that while the majority of the aptamers displays peak fluorescence intensity at around 1.5 hours to then decrease (Figure 2a, Figure S2a), two of the aptamers, namely tRNA-DF30-Pep and tRNA-1xPep, displayed a consistent increase in fluorescence signal to reach steady state intensity between four and ten hours (∼1400 f.u. and ∼3700 f.u., respectively) (Figure 2a, Figure S2a). Weak constitutive promoter J23118 resulted in early expression of aptamers (in the first 3 hours in DH10B) with exception of F30-4xPep and tRNA-1xPep in BL21, and F30-4xBroc in DH10B (Figure 2 b, Figure S2 b and c). Expression from pBAD_wt_ resulted in a low fluorescence signal of F30-1x, -4x and -8xPep within the first hour of induction and a stronger increasing signal from expression of tRNA-Broc, tRNA-enBroc and tRNA-DF30-Pep (Figure 2 c, Figure S2 b and c). Expression from pLux resulted in more consistent increase of fluorescence from all aptamers apart form F30-4xPep with a particularly strong increase from tRNA-enBroc in BL21 and tRNA- DF30-Pep in DH10B (Figure 2 d, Figure S2 b and c). While these different dynamic patterns can be explained by different aptamer or dye stability overtime in vivo, we reasoned they could also be explained by differences in growth profiles.

**Figure 2:**
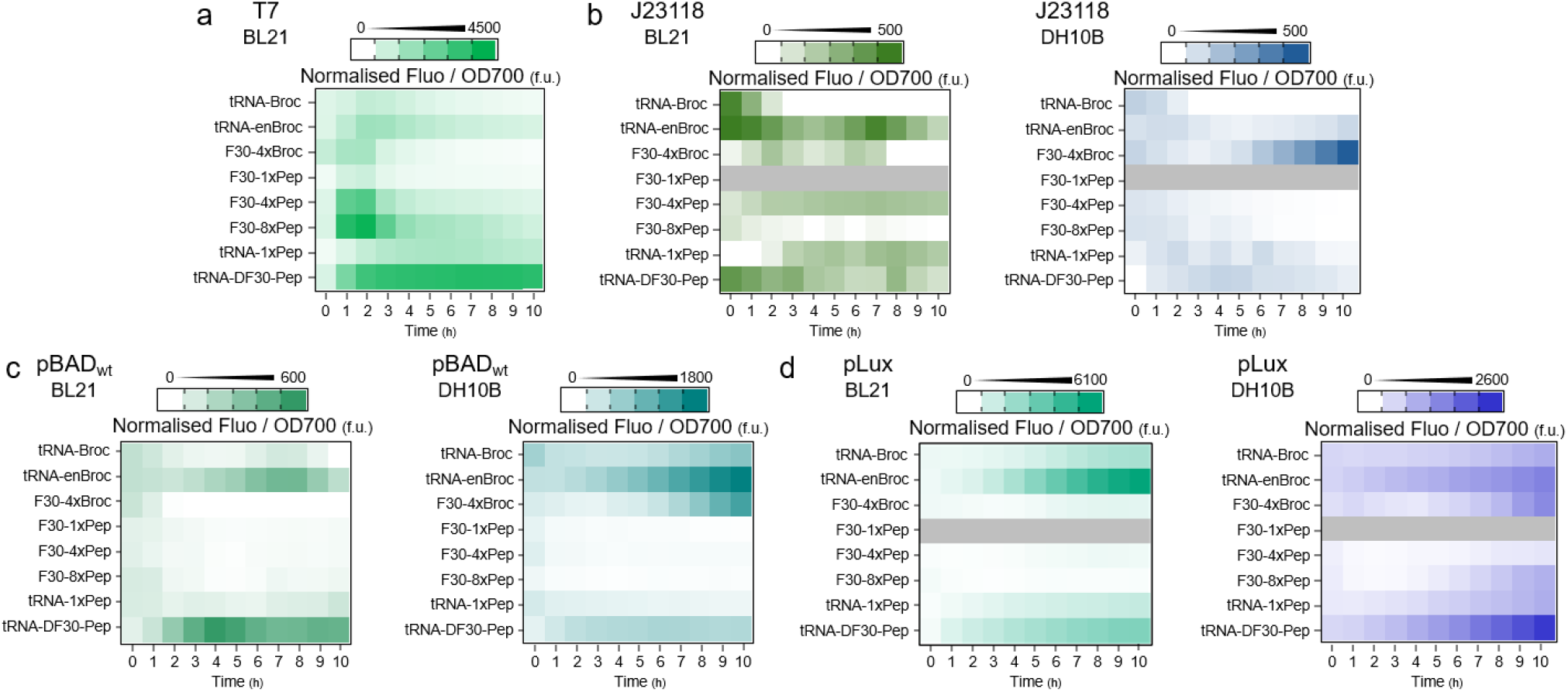
Dynamic performance over time of Pepper and Broccoli aptamers expressed in *E. coli*. Heat-maps of fluorescence levels per OD700 normalised by subtracting that of induced samples containing no fluorophore over a 10-hour window post induction. **a)** Performance of T7 constructs in BL21 induced with 1mM IPTG. **b)** Performance of J23118 constructs in BL21 (left) and DH10B (right). F30-1xPep aptamer performance is grey as it was not cloned under the J23118 promoter. **c)** Performance of pBAD_wt_ constructs in BL21 (left) and DH10B (right) induced with 0.2% arabinose. **d)** Performance of pLux constructs in BL21 (left) and DH10B (right) induced with 10uM HSL. F30-1xPep aptamer performance is grey as it was not cloned under the pLux promoter. (n=4)

We thus considered the growth profiles of the different samples. Interestingly, we observed that all aptamers expressed from T7 resulted in a delayed growth and lower carrying capacity (i.e. maximum OD reached by the culture) of BL21 compared to non-engineered cells, suggesting that their expression is causing burden in *E. coli* (Figure 3a). Different aptamers impacted growth differently with tRNA-DF30-Pep and tRNA-1xPep displaying the biggest impact (Figure 3a), leading to slower growth and corresponding slower increase of fluorescence over time. We observed a similar impact of aptamer expression on *E. coli* cell growth when aptamers were expressed from different promoters (Figure 3b-d) with the stronger impact observed for stronger promoters such as pLux (Figure 3d). tRNA-DF30-Pep displayed the biggest impact in both strains (Figure 3b-d) with the exception of pBAD_mut_ where tRNA-1xPep impacted growth the most (Figure S3).

**Figure 3:**
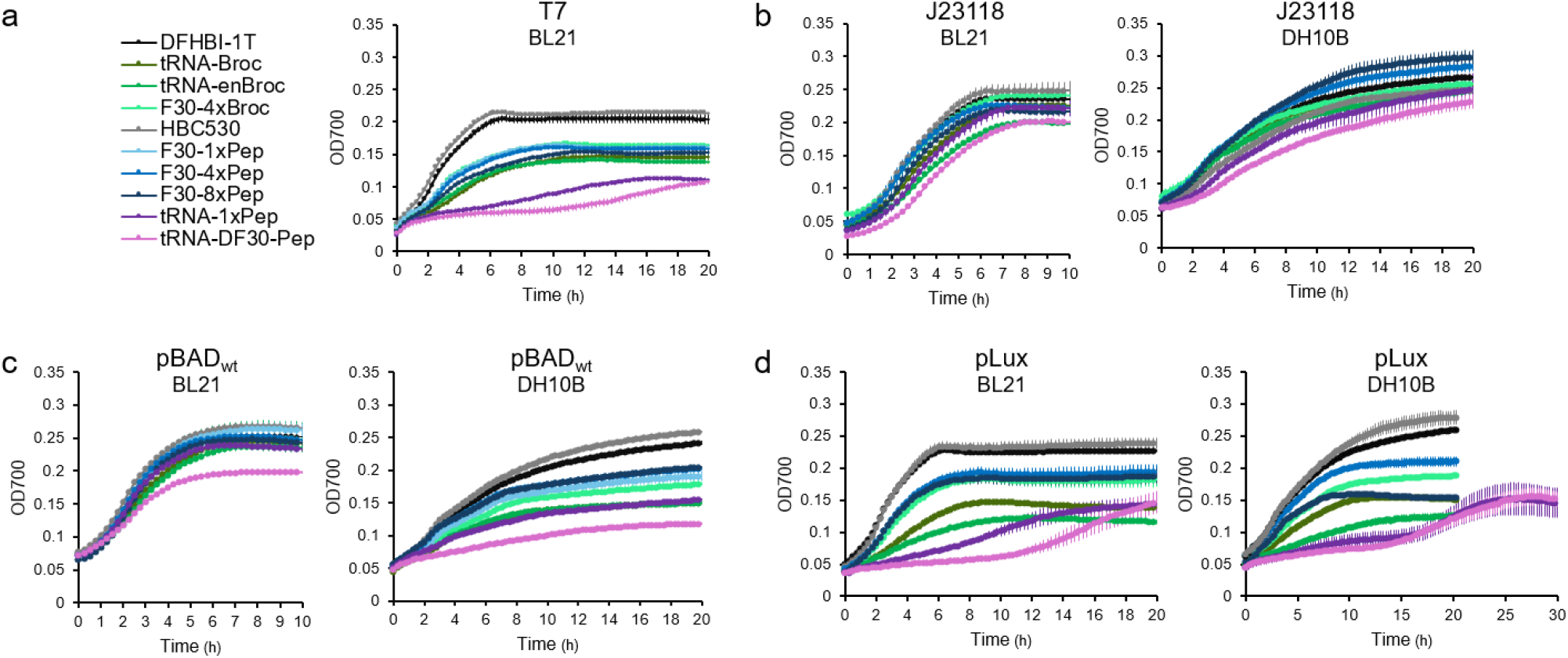
Impact of Pepper and Broccoli aptamers expression on their *E. coli* host. OD700 growth curves over time of *E. coli* cells supplemented with 160uM DFHBI-1T (grey) or 2uM HBC530 (black) and expressing Broc (green) or Pep (blue) aptamers from T7 in BL21 **(a)**, J23118 in BL21 (left) and DH10B (right) **(b)**, pBAD_wt_ in BL21 (left) and DH10B (right) **(c)**, or pLux in Bl21 (left) and DH10B (right) **(d)** following appropriate induction at t=0h. Data are presented as mean values +/- SEM (n=4).

Considering the secondary structures of the aptamers, we noticed that the observed aptamer-related burden appeared to be linked to the complexity of scaffolding (i.e. F30 < tRNA < tRNA-DF30) adopted within these RNAs, as exemplified in Figure S4a^16, 17, 34^. Increased scaffolding RNA complexity, thus appears to delay the time point at which maximal growth rate of engineered *E. coli* is found (Figure S4b).

To characterise the strength and functionality of the aptamers without the observed burden effect on the cells, we decided to test them *in vitro*. By adding the constructs to the PURExpress transcription and translation mixture, we followed fluorescence overtime by plate reader. At two hours post induction form a T7 promoter, F30-8xPep, tRNA-DF30-Pep and F30-4xPep were still the ones displaying higher fluorescence with 8.56-, 6.12- and 8.11-fold change compared to tRNA-Broc, respectively, but this time we noticed that all the Broc aptamers displayed lower expression compared to the Pep ones, with F30-1xPep being brighter than tRNA-enBroc (Figure S5a).

Given that many of the applications of synthetic biology entail adoption of *in vitro* systems for biosensing and prototyping^35^, we also sought to expand our *in vitro* characterisation to the commercially available Promega bacterial lysate. Generally, Pep aptamers performed better than Broc ones in this system, as we found with the PURExpress system (Figure S5 b-e), and we measured higher fold change expression than their live cell counterpart (e.g. tRNA-DF30-Pep expressed form J23118 reached 8.35-fold increase in fluorescence compared to tRNA-Broc in the lysate and 5-fold change when tested in DH10B) (Figure S5b). Interestingly, also for pBAD_mut_ the Pep aptamers reached up to 3.71-fold increase in fluorescence (F30-4xPep) behaving very differently when compared to tests in cells (Figure S5d, Figure S1d).

Overall, out of the seven tested aptamers, we selected tRNA-Broc and tRNA-enBroc, as these were the ones displaying the best combination of bright signal and burden, enabling maximisation of the readout and its early detection while imposing as low as possible impact on the host cells. These aptamers were thus cloned downstream of the htpG1 promoter on a low copy number plasmid and the constructs co-transformed with a previously characterised medium copy plasmid encoding for VioB-mCherry under the control of a pBAD_wt_ promoter^32^ known to cause burden in *E. coli* (Figure 4a). In this system, addition of the inducer arabinose leads to high expression of VioB-mCherry (Figure S6), and to a decrease in growth typical of a burdened phenotype (Figure 4b).

**Figure 4.**
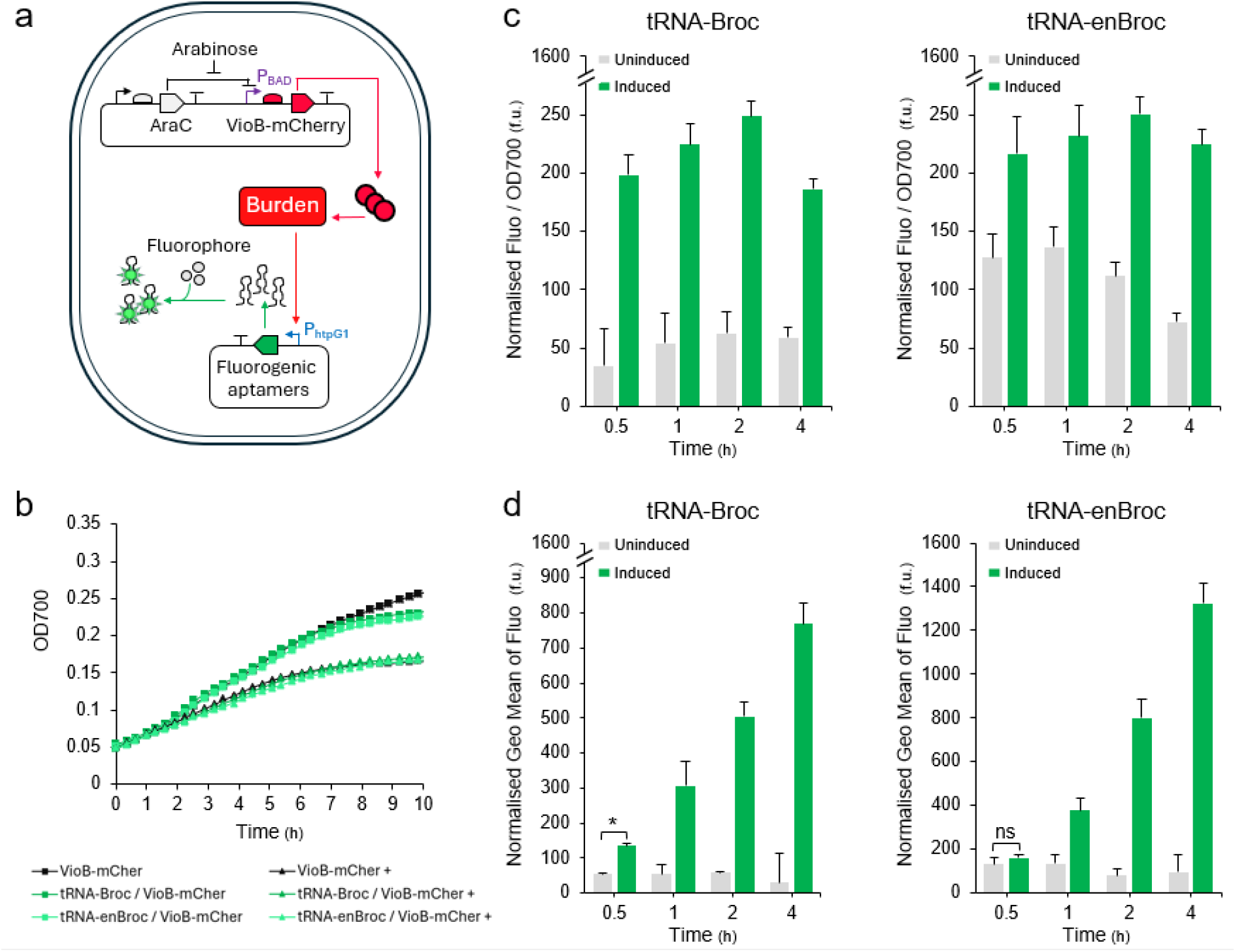
Monitoring cellular burden using fluorogenic aptamers in *E. coli*. **a)** Schematic of the constructs used in DH10B cells to induce burden and monitor it using fluorogenic aptamers. The expression of VioB-mCherry under pBAD_wt_ induces burden. The burden responsive promoter htpG1 subsequently triggers the expression of selected fluorogenic aptamers. **b)** OD700 growth curves over time of control DH10B transformed only with pBAD_wt_-VioB-mCherry plasmid (black) as well as DH10B co-transformed with pBAD_wt_-VioB-mCherry and phtpG1-tRNA-(en)Broc (green) plasmids. Cells were treated with 0% (square markers) or 1% (triangular markers) arabinose in M9 supplemented with 160uM DFHBI-1T. Data are presented as mean values +/- SEM. (=4) **c)** Bar chart resulting from plate reader measurements of sample fluorescence level per OD700 at 0.5, 1, 2 and 4 hours, normalised by subtracting that of non-engineered DH10B cells grown in same conditions with fluorophore. Uninduced samples (grey) and induced samples (green) were treated respectively with 0% and 1% arabinose at t=0h. Data are presented as mean values + SEM. (n=4) **d)** Flow cytometry resulting bar charts of geometric mean fluorescence at 0.5, 1, 2 and 4 hours, normalised by subtracting that of non-engineered DH10B cells grown in same conditions with fluorophore. Uninduced samples (grey) and induced samples (green) were treated respectively with 0% and 1% arabinose at t=0h. Data are presented as mean values + SEM (n=3). A two-sided t-test indicated that at 0.5 hours, fluorescence level in induced and uninduced samples are significantly different (*, 0.5>*p*>0.01) when using the tRNA-Broc biosensor, and non-significantly different (ns, *p* > 0.5) when using the tRNA-enBroc biosensor.

We tested the response of the two aptamers to burden first via plate reader assays, following fluorescence over time (Figure 4c). For all the aptamers tested, normalised fluorescence at 0.5, 1, 2 and 4 hours post induction in the induced samples was higher than in the uninduced control, thus confirming activation of htpG1. Interestingly, we could detect activation already at 30 minutes after heterologous expression induction, consistently with previous reports^3^. However, we noticed that the uninduced controls displayed high background fluorescence and that at 4h post induction activation of the sensor decreased compared to earlier time point, something we explained by considering cells are reaching late exponential phase. We thus decided to also confirm our results by flow cytometry at the same time points (Figure 4d). Flow cytometer measurements confirmed that induced samples activate transcription from htpG1, with tRNA-Broc displaying the highest signal-to-noise ratio from 30 minutes to 4 hours (2.44-, 5.63-, 8.35- and 25.77-fold change to uninduced at 0.5, 1, 2 and 4 hours respectively). tRNA-enBroc displayed brighter signal intensity (∼1325 f.u. at 4 hours post induction) but no clear activation at 30 minutes when compared to the uninduced controls (1.24- and 2.81-fold change to uninduced at 30 minutes and 1 hour respectively) (Figure 4d).

Overall, we identified two viable biosensor designs based on fluorogenic aptamers (tRNA-Broc and tRNA-enBroc) that can detect and measure burden in living *E. coli* cells in real time.

## Discussion

Cellular burden is a well-known hurdle for adoption of engineered organisms in biotechnology and synthetic biology applications. It limits host performance and overall yields. As of today, many tools have been developed to study and mitigate burden in bacteria. Here, we add to this collection by designing intracellular burden biosensors for the dynamic tracking of burden in living engineered *E. coli*. For these sensors, to be able to report on the burden response quickly, we selected fluorogenic RNA aptamers as the readout.

In biotechnology and cellular biology, fluorogenic RNA aptamers have gained interest for their suitability in live and dynamic RNA tracking. Nonoptimal stability, brightness and low signal-to-noise ratio are however among the limitations that have brought the community to develop many different RNA aptamers variants for their use in a range of organisms and applications.

To identify aptamers suitable for the design of our biosensor, we started by adopting plate reader-based measurements to characterise a library of seven aptamers of recent development, comprising variants of the Broccoli and Pepper aptamers, when these are expressed in two *E. coli* strains from a library of native and synthetic promoters. We first compared the brightness of the aptamer library when expressed in BL21 from the T7 promoter and then considered a library of different promoters to characterise the relative aptamer strength in BL21 and in DH10B cells, together with *in vitro* PURExpress and a different commercial bacterial lysate.

Our data highlight a context-dependent aptamer performance, with relative fluorescence strength being dependent not only on the aptamer sequence and secondary structure, but also on both the adopted promoter and cellular context. *In vitro* experiments also report differences in the relative performance of the aptamers compared to in vivo cell tests, highlighting how important it is to perform deep characterisation of aptamer performance across conditions and contexts, before selecting one for a given application.

We then considered the dynamic performance of the library and identified a scaffolding-related impact of different aptamers on cellular growth, with more burden observed for more complex scaffold structures. To the best of our knowledge, this is the first report of aptamer-related burden in *E. coli*. The reasons for such an impact are still unknown, and further experimental characterisation will be needed to understand how the scaffold interacts with the cellular host and the reasons for such a strong impact on the host. The observation that the choice of scaffold can not only impact the brightness of fluorogenic aptamers but also their impact on cellular physiology, calls for caution for those interested in adopting such systems for in vivo studies.

Finally, by selecting two aptamer variants with brighter signal but also lower burden, we developed a burden biosensor that triggers fluorescence in cells under the control of the burden-sensitive htpG1 promoter. Our experiments confirm that the htpG1 promoter can be used modularly with different aptamers to build burden biosensors and we demonstrate that these can be used as reporters to monitor the burden response of bacterial hosts to heterologous gene expression. Interestingly, the presence of the aptamer does not add to the burden imposed by VioB-mCherry alone, confirming that expression of a protein is still more burdensome for the cells than the sole RNA. htpG1 is also a native promoter likely weaker than the majority of the synthetic ones adopted in cell engineering, thus with minimal impact on the cell’s resources. One of the two biosensors, expressing tRNA-Broc, displayed a significant response within 30 minutes from induction of heterologous expression, consistently with our previous results reporting that the burden response downstream of htpG1 is upregulated already at 15 minutes after induction of heterologous protein expression^3^. Flow cytometry measurements appeared to yield better detection of the biosensor activation, i.e. less background fluorescence being displayed by the uninduced samples and consistent over time increase of the induced samples, likely due to plate reader measures capturing whole well fluorescence and thus also medium background signal.

Previously, biosensors were developed for mammalian systems and showed to be instrumental for tracking cellular stress in response to a variety of stimuli^36,37^. Our study develops a novel class of biosensors able to track burden live in *E. coli* cells. By adopting fluorogenic RNA aptamers as readout, these biosensors also ensure faster tracking of the burden response, compared to systems where fluorescent protein reporters are adopted.

In cell engineering, such burden-monitoring tools may be instrumental to aid identification of best performing construct designs with lower footprint on the host and better performance. We envision the biosensors developed here to be a useful resource to aid bioprocessing applications where growth and product yield maximisation are sought.

## Author contribution

AG and FC conceptualised the research and designed the experiments. AG, TC and CS performed experiments. AG, TC, CS and FC analysed the data. AG and FC wrote the manuscript. All authors read and edited the manuscript.

## Acknowledgements

The authors would like to thank Prof Hammond for providing the tD30-pepper aptamer sequence and Dr. Michael Booth for providing the enhanced-Broccoli aptamer sequence. This work was supported by the Biotechnology and Biological Sciences Research Council (grant BB/V00882X/1 to A.G and F.C.) and the EPSRC Centre for Doctoral Training in BioDesign Engineering (EP/S022856/1) (to TC, CS, YE, TE and FC). EF acknowledges support from BBSRC-NSF/BIO award 2020039.

## Methods

### Bacterial strains and DNA constructs

*E. coli* strains DH10B (K-12 F- λ-araD139 Δ(araA-leu)7697 Δ(lac)X74 galE15 galK16 galU hsdR2 relA rpsL150(StrR) spoT1 deoR ϕ80dlacZΔM15 endA1 nupG recA1 e14-mcrA Δ(mrr hsdRMS mcrBC)) and BL21(DE3) (F−ompT gal dcm lon hsdSB(rB−mB−) λ(DE3 [lacI lacUV5-T7p07 ind1 sam7 nin5]) [malB+]K-12(λS)) were used.

T7 constructs were cloned into pSCIB3 plasmid using inverse PCR to introduce the T7 promoter as a XbaI/NheI fragment and T7 terminator as BsaI/SpeI (KpnI/SpeI in the case of F30-1xPep). All fluorogenic aptamers were cloned downstream of T7 by annealing complementary ultramers generating overhangs of appropriated restriction sites.

For J23118 constructs, promoter sequences were cloned as a XbaI/PacI fragment with fluorogenic aptamers cloned as PacI/XhoI fragments upstream of a B0015_terminator sequence cloned as a XhoI/PstI fragment into the pSC1B3 plasmid.

pBAD_wt_ and pBAD_mut_ constructs were cloned into pSC1B3 by replacing VioB-mCherry from Ceroni et al^28^ with annealed fluorogenic aptamers cloned as NheI/SpeI fragments.

pLux constructs were cloned by inverse PCR of a pSC1B3 plasmid containing J23101-LuxR-pLux (a kind gift from Guy-Bart Stan)^37^ thus inserting PacI and NsiI restriction sites into which fluorogenic aptamers (PacI/PstI) were inserted.

All constructs were transformed both into *E. coli* strains BL21; J23118, pBAD and pLux constructs were also transformed into *E. coli* strains DH10B.

HtpG1 constructs, were cloned by replacing J23118 sequences from constructs in the pSC1B3 plasmid described above with htpG1 sequences as SfiI/PacI fragment upstream of the tRNA-Broc and tRNA-enBroc coding sequences.

Both htpG1 constructs were co-transformed into *E. coli* strains DH10B together with a pLys plasmid coding for expression of VioB-mCherry under the pBAD_wt_ promoter published in Ceroni et al^32^.

### Characterisation of RNA fluorogenic aptamer using plate reader assay

*E. coli* cells transformed with fluorogenic aptamer constructs were grown at 37 °C overnight with aeration in a shaking incubator in 500 ul of defined supplemented M9 media with the appropriate antibiotic. In the morning, 70 μl of each sample was diluted into 1 ml of fresh medium and grown at 37 °C with shaking for another hour (outgrowth). Then, 100 μl of each sample at approximately 0.05 OD700 was transferred into a 96-well plate (Greiner), inducers (1mM IPTG for T7 constructs, 0.2-1% arabinose for pBAD constructs and 10uM HSL for pLux constructs) were added and medium was supplemented or not with the correct fluorophore (160µM DFHBI-1T for Broc variants and 2µM HBC530 for Pep variants). The plate was sealed with BreathEasy and place into a Tecan plate reader, incubated at 37 °C with orbital shaking at 1,000 rpm for up to 20h, performing measurements of OD (700nm) and fluorogenic aptamer fluorescence (λ_ex_=485nm; λ_em_=530nm) every 15 minutes. Measurement of biosensors activation following induction of VioB-mCherry expression was performed as described above with measurements every 15 minutes of OD (700nm), fluorogenic aptamer fluorescence (λ_ex_=485nm; λ_em_=530nm) and mCherry fluorescence (l_ex_=587nm; λ_em_=610nm).

### Characterisation of fluorogenic RNA aptamers using bulk *in vitro* transcription-translation reactions

Plasmids encoding the relevant aptamer design were prepared from bacterial cultures. For the constructs under the control of the T7 promoter, expression was performed using the PURExpress® In Vitro Protein Synthesis Kit (NEB). For the non-T7 constructs, expression was performed using the *E. coli* S30 Extract System for Circular DNA (Promega). All reactions were treated with murine RNase inhibitor to prevent RNA degradation. Reactions were assembled in 384-well plates with either DFHBI-1T or HBC530 fluorophore at a final concentration of 160 µM or 2 µM, respectively. For constructs expressed from arabinose-inducible promoters, wells were supplemented with 1% arabinose. For constructs expressed from LuxR-inducible promoters, wells were supplemented with 10 µM 3-oxo-hexanoyl homoserine lactone (HSL). Fluorescence intensity was measured in a ClarioSTAR multi-mode plate reader for 4 hours at 37°C, shaking at 250 rpm.

### Characterisation of RNA fluorogenic aptamers using flow cytometry

Glycerol stocks of the DH10B cells containing the plasmid encoding the relevant aptamer design, with or without an integrated arabinose induced mCherry expression cassette, were spread on LB agar plates containing 50 µg/mL Chloramphenicol +/- 50 µg/mL Ampicillin and incubated overnight at 37°C. Colonies were picked and inoculated in 500 µL M9 media containing appropriate antibiotics before being incubated at 37°C, shaking at 250rpm, for 16 hours. 140 µL of overnight cell culture was diluted into 2 mL M9 with antibiotics and incubated for 1 hour, at 37°C, shaking at 250 rpm. 100 µL of each outgrowth culture were transferred to a 96-well plate, 160 µM DFHBI-1T fluorophore was added, and samples were treated with 0% or 1% arabinose, before being incubated at 37°C, shaking at 250 rpm. At 0.5, 1, 2 and 4 hours after induction, 20 µL of these cell cultures were transferred to 180 µL PBS in a fresh 96-well plate. Fluorescence was then measured on the Attune CytPix, Invitrogen, using the CytKick Max autosampler, Invitrogen. Data analysis was performed on FlowJo v10 software.

## Supplementary information

**Figure S1.**
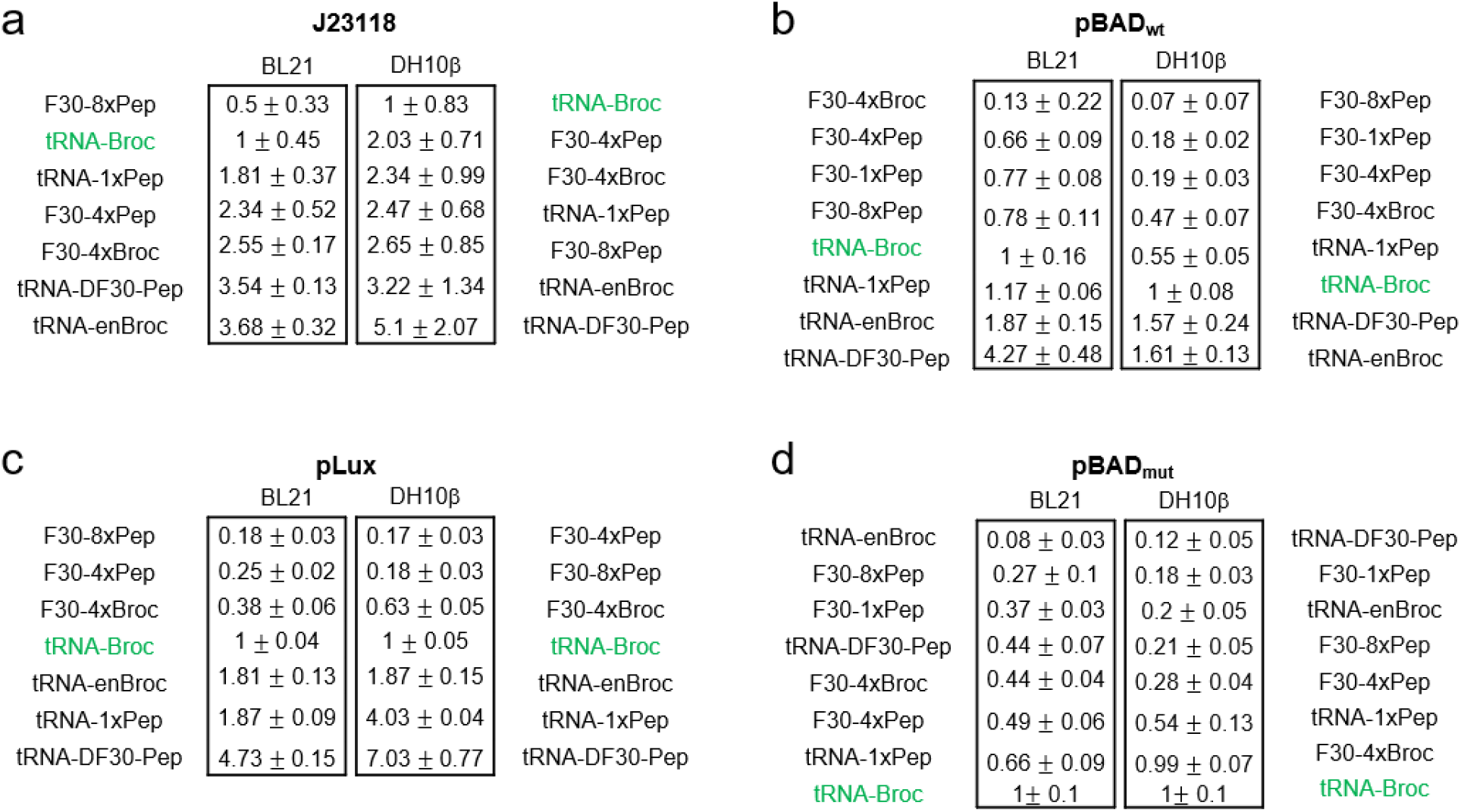
Tables of RNA aptamer fluorescence fold change to tRNA-Broc at maximum growth rate. Tables of fluorescence fold change +/- SEM in comparison to that of tRNA-Broc (green) at maximum growth rate, when fluorogenic aptamers are expressed either in BL21 or DH10B under J23118 **(a)**, pBAD_wt_ **(b)**, pLux **(c)** or pBAD_mut_ **(d)** (n=4).

**Figure S2.**
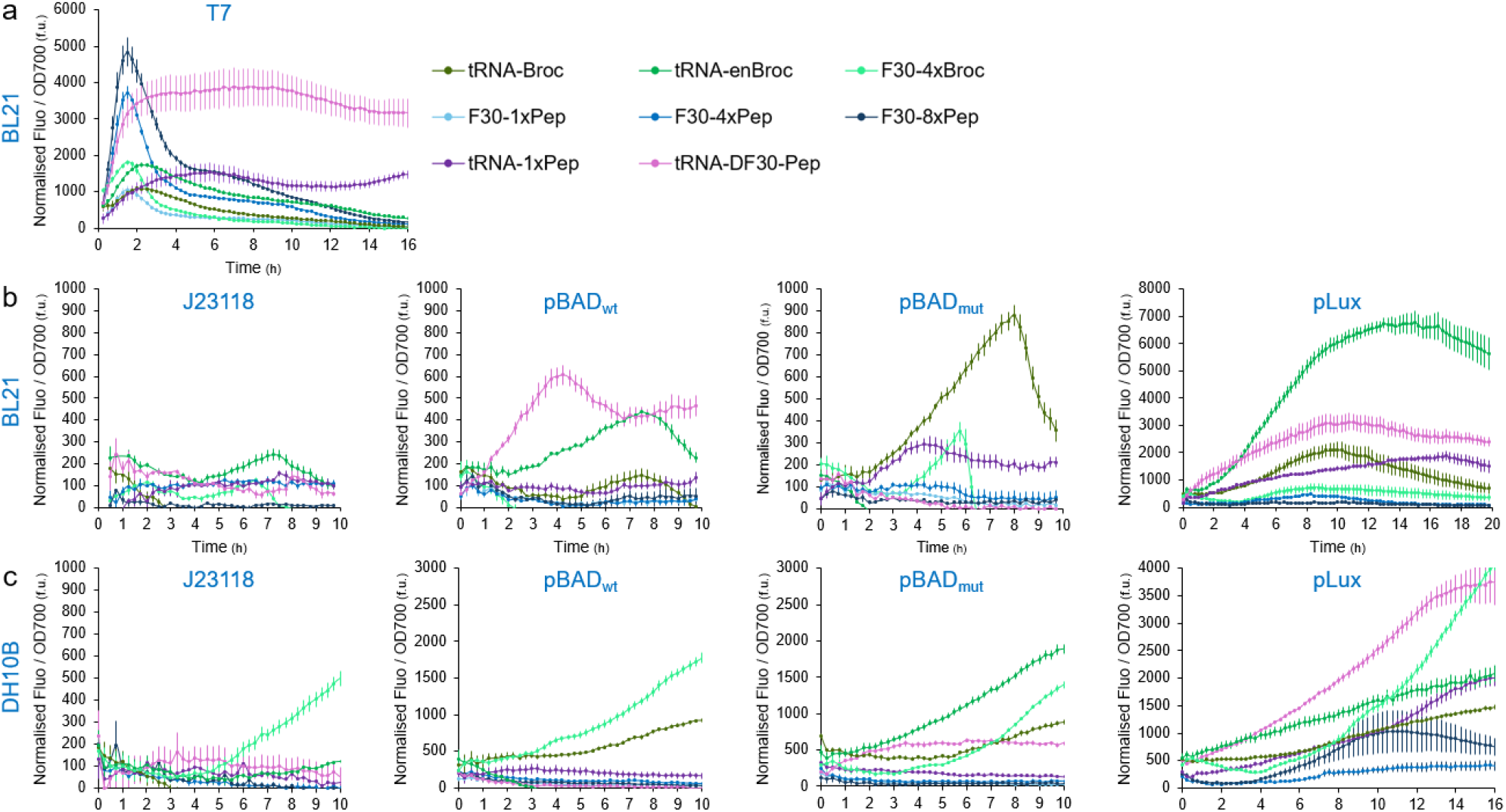
Normalised Fluorescence / OD700 in BL21 and DH10B. Fluorescence level per OD700 over time of induced samples supplied with correct fluorophores (160uM DFHBI-1T or 2uM HBC530), normalised by subtracting that of induced samples lacking fluorophores. Resulting normalised fluorescence per OD level over time of Broc aptamer variants (green), F30 Pep aptamer variants (blue) and tRNA Pep aptamer variants (purple) expressed under T7 promoter in BL21 are plotted in **(a)**, while their normalised fluorescence per OD when expressed under J23118, pBAD_wt_, pBAD_mut_ and pLux are plotted in **(b)** for BL21 and **(c)** for DH10B. Expression from T7 promoter was induced by 1mM IPTG, while expression from pBAD promoters was induced by 0.2% arabinose and expression from pLux promoter was induced by 10uM HSL added at t=0h. Data are presented as mean values +/- SEM (n=4).

**Figure S3.**
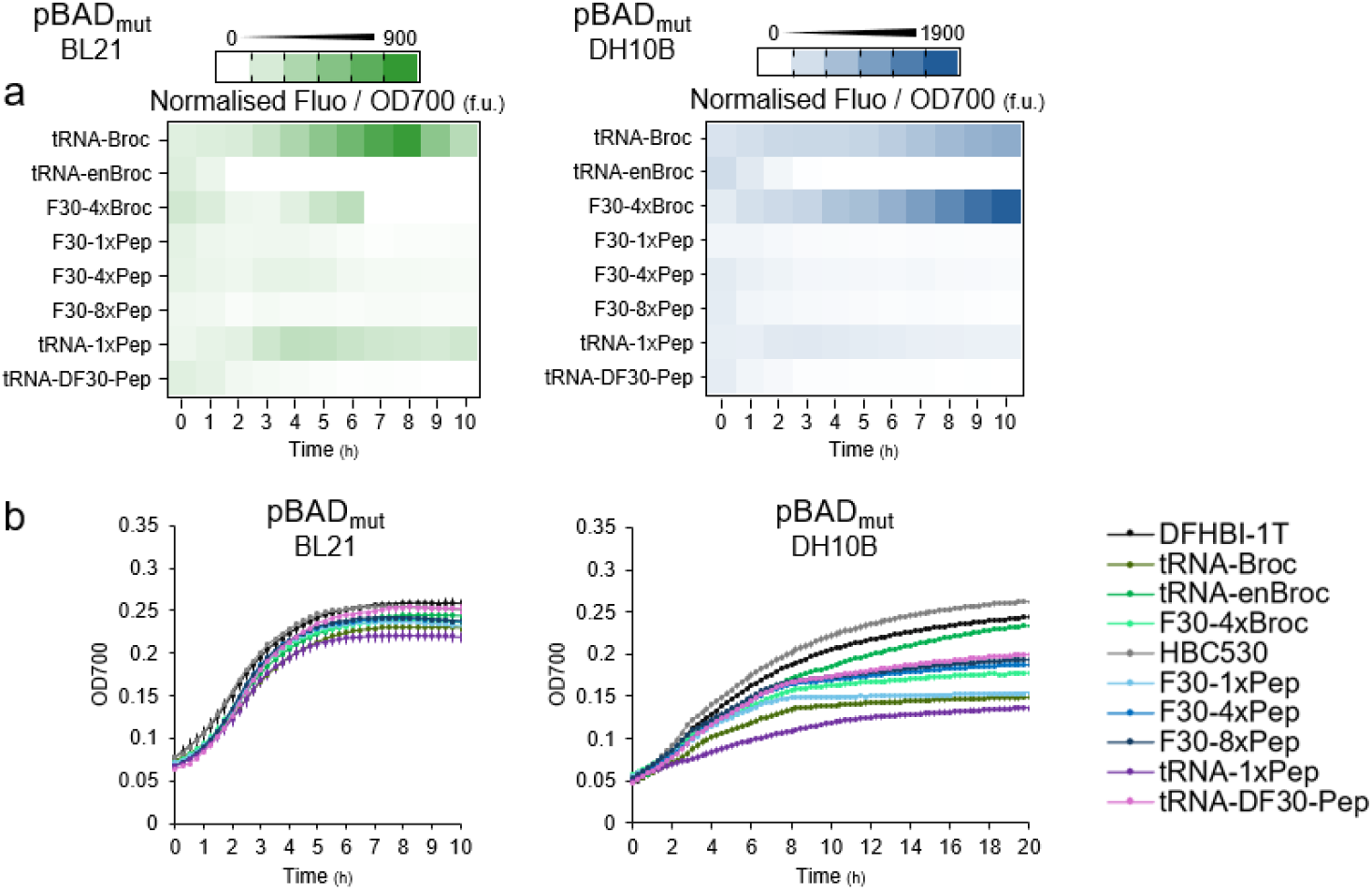
Dynamic performance of aptamers expressed in E. coli from the pBAD_mut_. **a)** Heat-maps of fluorescence levels per OD700 normalised by subtracting that of induced samples containing no fluorophore over a 10-hour window post induction. Performance of pBAD_mut_ constructs in BL21 (left) and DH10B (right) induced with 0.2% arabinose at t0h. (n=4) **b)** OD700 growth curves over time of *E. coli* cells supplemented with 160uM DFHBI-1T (grey) or 2uM HBC530 (black) and expressing Broc (green) or Pep (blue and purple) aptamers from pBAD_mut_ in BL21 (left) and DH10B (right) following 0.2% arabinose induction at t=0h. Data are presented as mean values +/- SEM (n=4).

**Figure S4.**
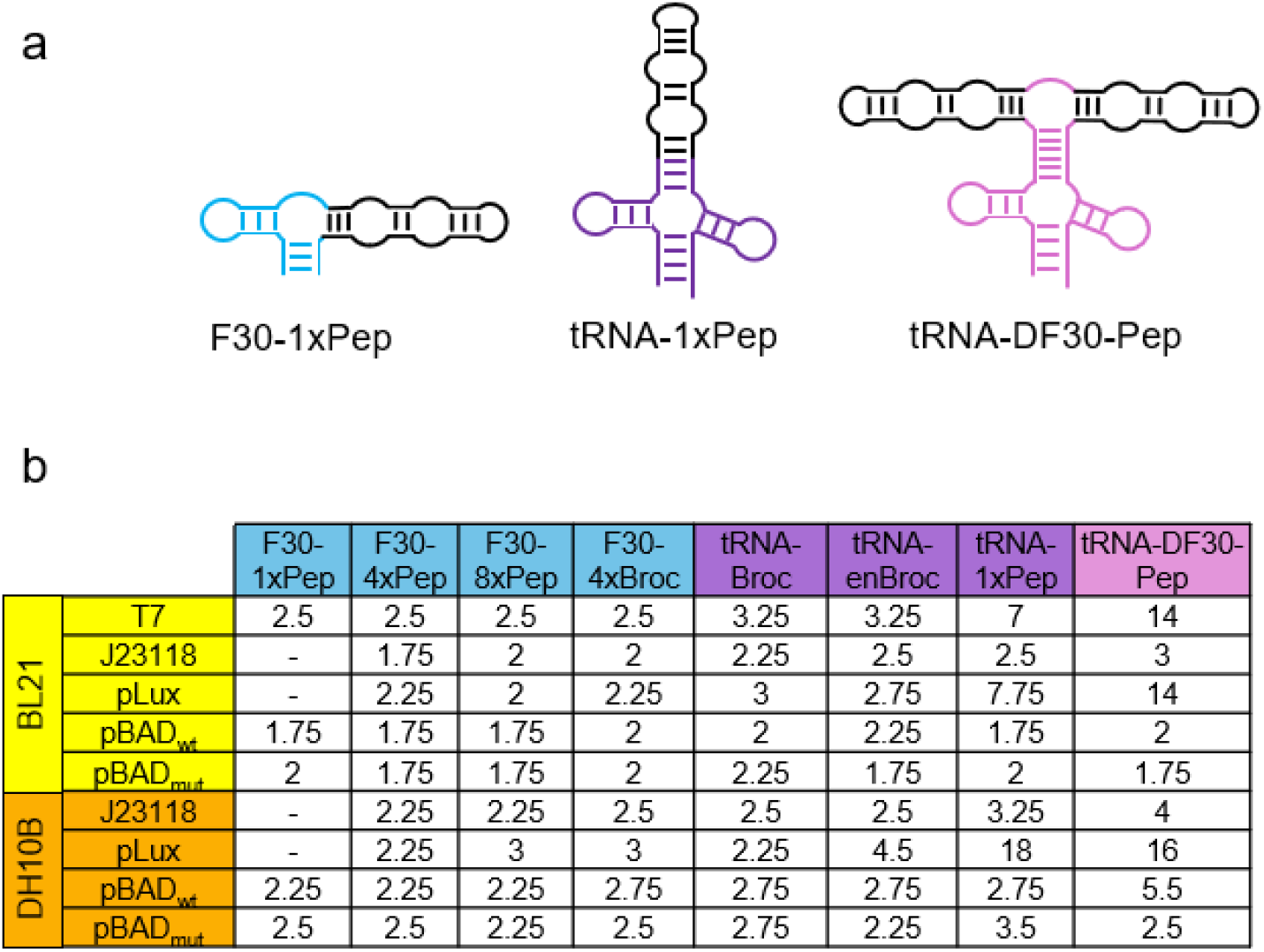
Impact of scaffolding RNA on fluorogenic aptamer cellular burden. **a)** Schematics of F30 (blue), tRNA (dark purple) and tRNA-DF30 (light purple) RNA scaffolds linked to Pep (black). **b)** Table of *E. coli* BL21 (yellow) and DH10B (orange) cells maximum growth rate time points when expressing fluorogenic aptamers linked to F30 (blue), tRNA (dark purple) or tRNA-DF30 (light purple) (n=4).

**Figure S5.**
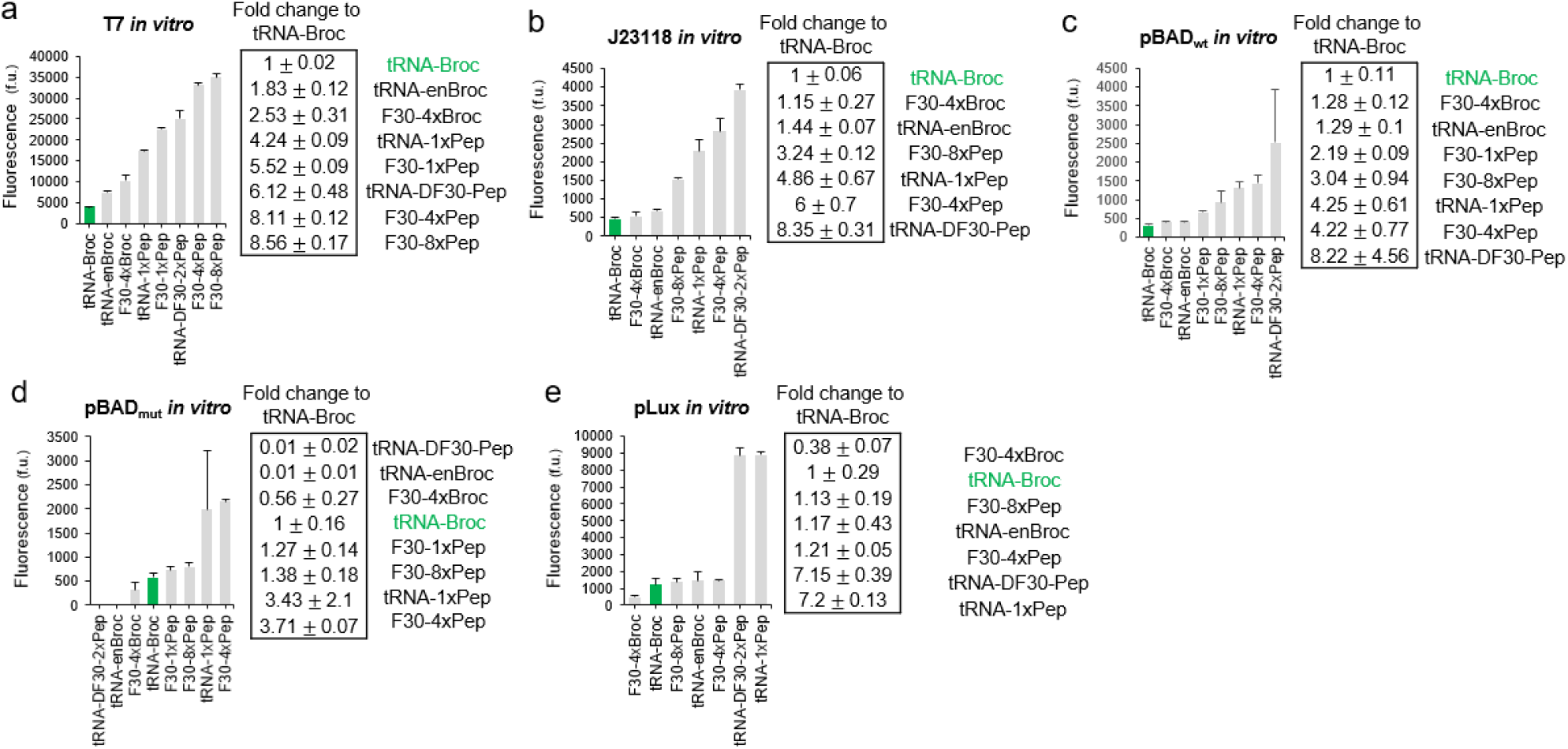
In vitro characterisation of fluorogenic aptamers expression under T7, J23118, pBAD_wt_, pBAD_mut_ and pLux. *In vitro e*xpression levels of T7 **(a)**, J233118 **(b)**, pBAD_wt_ **(c)**, pBAD_mut_ **(d)** and pLux **(e)** constructs with a bar chart of fluorescence levels at two hours post reaction start (left) and resulting table of fluorescence fold change in comparison to tRNA-Broc (right). T7 constructs were expressed in PURExpress transcription and translation mixture, while J23118, pBAD_wt_, pBAD_mut_ and pLux were expressed using Promega bacterial lysate. Standard tRNA-Broc sample is highlighted in blue. Data are presented as mean values +/- SEM (n=2).

**Figure S6.**
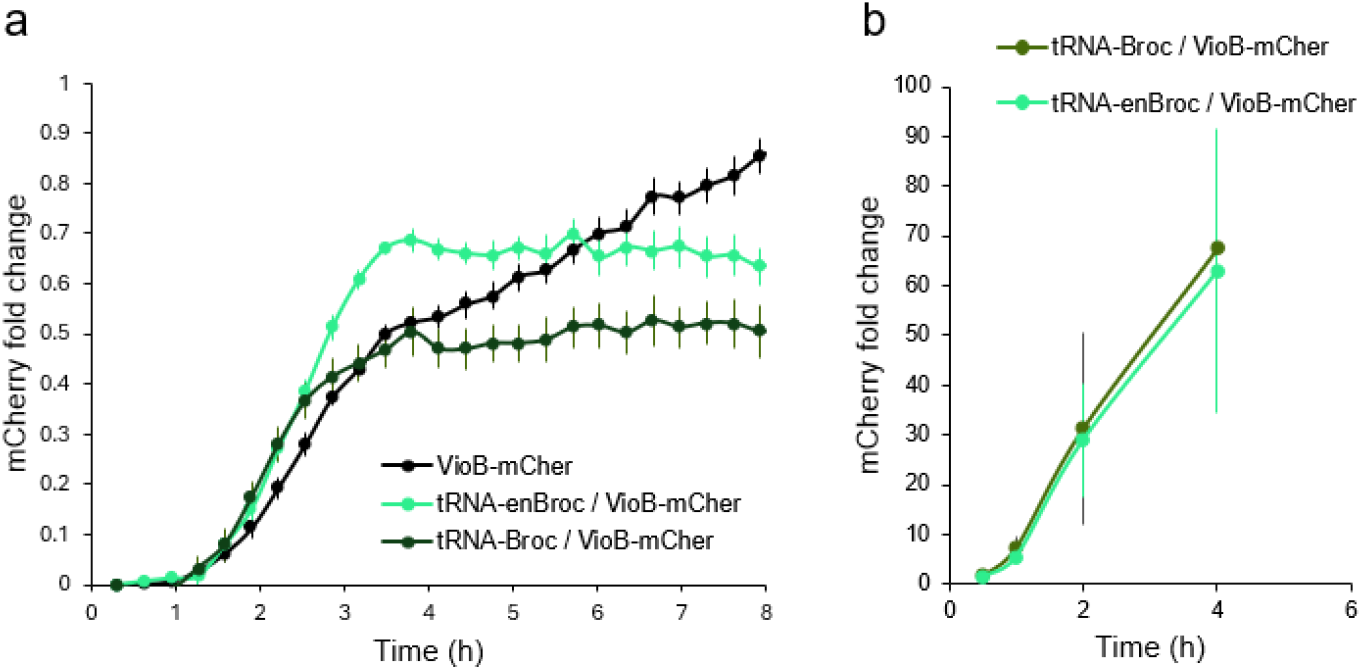
Fold change of induced VioB-mCherry fluorescence. **a) Over time** fold change in mCherry fluorescence on OD700 for induced samples (treated with 1% arabinose) normalised over uninduced samples (treated with 0% arabinose) as measured in a plate reader. Data are presented as mean values +/- SEM (n=4). **b)** Over time fold change in mCherry geometric mean fluorescence for induced samples (treated with 1% arabinose) normalised over uninduced samples (treated with 0% arabinose) as measured by flow cytometry. Data are presented as mean values +/- SEM (n=3).

